# Extended high frequency hearing influences cortical response amplitudes to speech

**DOI:** 10.64898/2026.05.23.727071

**Authors:** Rodrigo Donoso-San Martin, Stefan Fink, Konrad Dapper, Etienne Gaudrain, Deniz Başkent, Sarah Verhulst, Csaba Harasztosi, Wibke Singer, Manuel S. Malmierca, Markus Siegel, Ernst Dalhoff, Stephan M. Wolpert, Christoph Braun, Lukas Rüttiger, Marlies Knipper

## Abstract

Since young adults hear sounds up to 20 kHz, the loss of extended high-frequency hearing (**EHF**; above 8 kHz) is a hallmark of age-related hearing loss, often progressing from early lifetime. However, this deficit frequently goes undetected because routine clinical hearing tests and most hearing aids are currently limited mostly up to 8 kHz. EHF hearing has been linked to deficits in speech perception in noise and to self-reported hearing. However, it remains elusive how EHF hearing influences speech intelligibility. Here we recorded neuromagnetic brain responses using magnetoencephalography (MEG) within a frequency-tagging speech paradigm designed to probe hierarchical levels of attention and memory-dependent speech processing and recognition. Auditory evoked cortical magnetic field (AEF) responses were significantly reduced in both left and right brain hemispheres in individuals with impaired EHF hearing compared to those subjects with rather preserved EHF hearing. A gradual reinforcement of left-hemispheric AEF seen over age was not observed in young adults (19-29 y) with preserved EHF hearing. This was linked to stronger auditory brainstem responses (ABR), reflecting better neural synchronized auditory responses at stimulus onset. The reinforced left hemispheric dominance in young adults with impaired EHF hearing, in contrast, was linked to lower ABRs. Our findings suggest that sound energy above 8 kHz contributes through its impact on stimulus-onset synchrony to phase locking of oscillations in the auditory cortex to intelligible speech. Together, the results highlight the need to reconsider the neglect of EHF hearing in both audiological assessment and hearing aid design.

**Significance:** We show here that deficits in extended high-frequency **(EHF)** hearing, up to now neglected in routine clinical audiometry and hearing aid technology, lead to reduced cortical evoked auditory field **(AEF)** response amplitudes to attended and unattended speech, even at a young age. A gradual increase in reinforced left-hemispheric AEF responses during attended speech does not occur in young people with good EHF hearing; this is linked to better synchronization of neural responses at the onset of sound. This suggests a crucial role of sounds containing energy above 8kHz in minimizing the need for cognitive resources during active listening. Collectively, our results challenge current clinical practices and underscore the need to incorporate EHF hearing into audiological assessment and hearing aid design.

## Introduction

Age-dependent hearing impairment has recently been identified as a major modifiable risk factor for cognitive decline in later life (1–3). A growing body of evidence has linked hearing loss and dementia to a deterioration of brain resources due to an impoverished acoustic environment and to the reduced availability of cognitive resources that support active listening in difficult conditions (4, 5). Active listening and speech intelligibility require cognitive resources, including the hippocampus, that, via numerous efferent auditory connections from the medial temporal lobe enhance the phase coupling of slow brain oscillations in conjunction with auditory input activity (6–8). An increased reliance on these cognitive resources, for example due to impaired hearing, could – as has been suggested – become a risk factor for earlier-onset dementia, for example by recruiting increased activity of the medial temporal lobe when listening in the real world (5, 8–10).

Keeping this in mind, previous studies suggesting a role of extended high frequency (EHF) hearing above 8 kHz for speech recognition and speech perception in noise (11, 12) should be taken into account. Speech in noise - due to its particular greater cognitive demands displays a special relationship to cognitive performance and the development of dementia (4). Unfortunately, to date neither diagnostic audiometry nor hearing restoration through hearing aids cover frequencies above 8 kHz, although young healthy adults can hear tones up to at least 20 kHz (13). Our previous studies have suggested that EHF between 8-16 kHz may influence the auditory transfer function at stimulus onset by improving synchronized responses and modulating low-frequency hearing (14, 15). Here, we ask how EHF might influence cortical activity during speech intelligibility. Speech comprehension during active listening depends critically on the integration of auditory speech cues with higher-order (non-sensory) linguistic elements that drive speech intelligibility during a so-called cerebro-acoustic process (7, 16). During this process, phase locking at brain oscillatory frequencies 4 to 7 Hz between auditory and cognitive resources exhibits maximal power (6, 7, 17). We therefore presented and tagged sentences within 4 Hz rates (4-word sentence) and 3.75 Hz rates (5-word sentence) to substantiate the reproducibility of observed altered cortical AEF.

To ensure that our paradigm effectively capture intracortical activity changes during speech comprehension, we additionally presented the 4-/5-word sentences by a ‘target’ speaker both in isolation and in the presence of a ‘distracting’ second speaker: A distractor only exhibits minor attenuation of signal strength for neural signals linked to a higher hierarchical speech percept (6, 18). Using high-resolution 275-channel MEG recording in young, middle-aged, and older subjects, we related entrained cortical responses to 4-/5-word sentences were related to detailed audiometric measurements, including information about EHF hearing and stimulus-onset precision that were previously obtained in the same subjects (14, 15). This approach allowed us to obtain initial insight into how EHF hearing influence auditory or cognitive processes during active language comprehension.

## Methods

### Participants

The consent form and experimental procedures were approved by the ethics committee of Tübingen University (Faculty of Medicine; ethical approval number 392/2021BO2). Written informed consent was given by all participants. All methods followed the Declaration of Helsinki by the World Medical Association (WMA) for human research ethics (14, 15).

Eighty-one participants (18 to 76 years, mean 47 ± 16) were recruited, from which thirteen participants were excluded because of motion and technical artifacts contaminating the physiological recordings. All gave written informed consent. The remaining 68 subjects were subdivided in 3 groups of young, middle-aged and older subjects and 2 groups having good or impaired EHF hearing (See SI appendix, Table S1). Whether participants fell into the preserved (here named as ‘good’) or impaired EHF group was determined based on the regression line of the EHF threshold against age. Participants whose EHF threshold was below the age-typical level were assigned to the good EHF group, whilst those whose EHF threshold was above the age-typical level were assigned to the impaired EHF group.

### Pure-tone audiometry and ABR measurements

Audiometric measurements (pure-tone audiometry and ABR measurements) were previously described in detail (14, 15). In brief: The pure-tone threshold was measured using an AT1000 Audiometer (Auritec, medizindiagnostische Geräte GmbH, Hamburg, Germany). The default pure-tone audiometric thresholds from 0.125 to 10 kHz, and the UCL (0.25, 0.5, 1, 2, 4, and 6 kHz) were measured using Beyerdynamic AT1350A on-ear headphones (Beyerdynamic, Heilbronn, Germany). In addition, EHF thresholds were measured using Sennheiser HDA300Pro (Sennheiser, Wedemark-Wennebostel, Germany) on-ear headphones at the frequencies 11.2, 12.5, 14, and 16 kHz. The Sennheiser HDA300 Pro achieves a nominal level of 123 dB SPL (6 Hz to 23 kHz). Pure-tone average thresholds for extended high frequencies (11.3, 12.5, 14, and 16 kHz) were derived from the right-ear thresholds. The ABR amplitudes (Fig. 3, 4, ABR wave I-VI), were measured monaurally using three electrodes (Neuroline 720, Ambu, Bad Nauheim, Germany), and the actiCHamp Plus64 amplifier as described in (14). Stimuli were generated using a Scarlet Focusrite 8i8 gen 3 soundcard (Focusrite, Buckinghamshire, UK) and presented through ER2 transducers and disposable ER1-14A earpieces (Etymotic Research, Elk Grove Village, IL, USA). The ER2 are limited to approximately 8 kHz.

### MEG data recording

Data were collected using a CTF Squid MEG System (OMEGA 275). The participants were seated in an upright position and instructed to look at a fixation cross in front during the entire measurement. They were instructed not to suppress blinking but to blink naturally. Participants were constantly monitored with a camera and microphone. Auditory stimulation was presented via ERA tone 3A 300 Ohm transducers (Etymotic Research) attached to tubes of 2 m length. Participants responded by pressing a confirm button interrupting the light beam between optical fibers.

### MEG task and stimuli

The task was designed to explore how attention and distraction affect brain responses during speech processing. The masking of an attended speaker by a second speaker was chosen to replicate the cocktail-party effect, often experienced by older individuals and possibly related to presbycusis (19). In a concurrent speech stimulus (SI appendix, Fig. S1) frequency-tagged brain responses were analyzed in the speech rhythm of a target speaker with and without a second speaker present. Speaker’s voices were chosen from different genders to ensure that they sounded sufficiently distinct. The participants were instructed to pay attention to one of the two concurrent voices (herein after referred to as “attended”) to recognize a target sentence that was presented visually on a screen in front of them (SI appendix, Fig. S1). After they had read the sentence on the screen, the participant could start the auditory stimulation by button press (SI appendix, Fig. S1).

During each recording session, four auditory stimulation blocks were presented. The first three blocks (I-III) focused on studying speech comprehension, while the last block involved recording the brain activity at rest. Each block was divided into 40 segments of 30 seconds, at the beginning of which the target sentence was displayed on the screen in front of the participant (SI appendix, Table S2). Each segment consisted either of 29 4-word or 23 5-word sentences. In the 4-word sentence condition, words were presented at a rate of 4 Hz leading to a sentence rate of 1 Hz, in the 5-word sentence condition the word rate was 3.75 Hz and the sentence rate 0.75 Hz (SI appendix, Table S2). The choice of the different frequencies minimized the crossover of harmonic word or sentence rates. The target sentence was presented either three times or twice per segment, depending on the sentence rate. Participants were asked to indicate each target sentence by a button press.

To exclude participants memorizing single words of the target sentence without listening to the whole sentence, the speakers used all words several times in different sentences. The instruction was to follow the target speaker and to ignore the second speaker. In the two-speaker conditions, the target speaker spoke the first sentence (the target sentence) before the distractor speaker started. The participants were instructed to push the button whenever they identified the target sentences (SI appendix, Fig. S1). Frequency tagging of evoked power in fT^2^/Hz at the sensor level was used to analyze the cortical responses to either 4- or 5-word rates.

### Stimulus design

The sentences used as stimuli were based on natural speech recorded in a soundproofed chamber. The speakers produced the words while listening to a metronome, in order to keep the syllable pace of 4 or 3.75 Hz. Any deviations from the rhythm were digitally corrected using Audacity 3.1 (Audacity Team, Free Software Foundation Inc, https://fsf.org), by equalizing the duration of the individual word snippets without changing pitch (*Subband Sinusoidal Modeling Synthesis*). During the experiment, the volume of the stimulation was varied until the subjects reported that it corresponded to the volume they would choose on their phone while listening to a person speaking in a quiet environment. This ensured a comfortable and sufficient sound level for each subject.

### MEG data preprocessing

MEG signals were preprocessed using second-order gradiometer correction. Next, two different bandpass filters (FieldTrip third-order ‘but’ filter, and bidirectional, third Order) were applied to the continuous data; one with 0.1 to 100 Hz and the second with 0.1 to 40 Hz. The latter filter was used prior to computing the independent-component-analysis **(ICA).** After filtering, data were epoched and down-sampled from 1172 Hz to 512 Hz. Filtered data were visually inspected to exclude any epoch or channels with particularly large amplitude variance. Subjects with more than 10 % of their epochs removed were excluded from further analysis, resulting in N=68 participants remaining for the data analysis. **Fast-ICA** using the ‘runica’ method was applied to remove noise from the MEG recordings. ICA was computed independently for each of the four measurement blocks. The resulting ICA components were visually inspected to remove eye movement, heartbeat, and muscle artifacts. The visual inspection was conducted without considering the participants’ age or other biomarkers. Subsequently, a fast Fourier transform was performed using the ‘mtmfft’ (Multi-Taper Method Fast Fourier Transform) for time-frequency analysis with a Hann window taper to isolate the different frequency-tagged responses.

The brain responses tagged by frequency to either 4 Hz (4-word sentences) or 3.75 Hz (5-word sentences) were analyzed from the first three blocks as four different stimulus conditions: (i) one-speaker attended with only the target speaker present and the participant paying attention to this speaker; (ii) two-speaker attended, with one speaker to attend to and a distracting second speaker to ignore, (iii) two-speaker unattended, taking into account the responses synchronized with the distracting speaker’s rhythm, i.e. the unattended speech stream, (iv) control as quantified crosstalk between word frequencies, defined by brain responses at 3.75 Hz and 4 Hz for a single speaker producing words at 4 Hz and 3.75 Hz, respectively.

### Evoked sensor-level power at a given frequency

For each subject and condition, the evoked power was calculated independently for the male and female speaker by first averaging all trials of each condition and computing the square of the modulus of the Fourier coefficients. To evaluate the quality of the recording, the evoked power’s noise level at a given frequency and condition was estimated by randomly shuffling the phases of the Fourier components before coherent averaging and sequence computation of the power of the noise. This shuffling procedure was repeated a thousand times for each subject. The average of all permutations estimates the noise level at a given frequency and was used to compute the signal to noise ratio **(SNR)** for each channel, frequency, and participant. These SNR values were then averaged across participants. Finally, the responses from the male and female speakers were averaged. The result represents a conservative estimate of the statistical significance of the recorded evoked activity for each sensor at a specific frequency.

### The time course of the absolute value of neuromagnetic responses

The time course of the event-related fields was obtained by averaging the absolute magnetic field amplitudes across trials and sensors, separately for the left and right brain hemisphere., To avoid the cancellation of dipolar neuromagnetic brain signals when averaging across sensors of one hemisphere, signal amplitudes were rectified at individual sensors.

### Statistical testing

**The sensor power-level** difference between conditions at a specific frequency was assessed using comprehensive permutation analysis. Therefore, the L^2^ norm of the difference of two conditions was compared with the difference of random permutations of the condition labels (’one speaker’, ‘two speaker’, ‘distracting speaker’, ‘control’). The L^2^-norm in this context refers to the average, calculated across the channels, of the square of the difference of evoked power. To eliminate the influence of inter-subject variability, all permutations were achieved within each subject. Statistical analysis was performed twice, once with the raw data and once with the data rescaled by their average power. The latter tested topographic distribution, rather than the total MEG-Power. Each time, 30,000 permutations were computed.

For the **time course of the absolute value of the neuromagnetic strength,** event-related fields were calculated from all trials of one of the four conditions centered on the rhythm of the word rate. The absolute values across all channels of one hemisphere for one condition were averaged. From the resulting time course of 1 s and 1.33 s for 4-word and 5-word sentences, respectively, the time course of the control condition averaged for all participants within the compared groups was subtracted from the one-speaker, two-speaker attended, and two-speaker unattended data. The resulting normalized time course of the response was inspected for peak response amplitudes within time windows defined by the word rhythm (0.05-0.25, 0.25-0.5, 0.5-0.75 and 0.75-0.95 s for the 4-word rhythm and 0.05-0.266,

0.266-0.533, 0.533-0.8, 0.8-1.066, and 1.066-1.333 for the 5-word rhythm). Average-peak event-related fields were statistically compared for the factors hemisphere, word-rate, and stimulus condition by 3-way ANOVA for repeated measurements (matching factors hemisphere and word rate) and tested by pairwise comparison using Sidak’s multiple comparisons test at significance level alpha=0.05 (GraphPad Prism 10.3.1, GraphPad Software, Boston MA, USA). Since neither word rate itself nor its interaction with hemisphere and stimulus condition significantly contributed to the total variance (p>0.05), word rate was not considered as a factor in subsequent ANOVA comparisons for EHF hearing (good versus impaired) and age (young, middle aged, old). Subsequent statistical comparisons were performed using 2-way repeated-measurement ANOVA (matched for the factor hemisphere) considering the factors EHF hearing (good and impaired) and hemisphere (left and right). Because the data for the participant groups separated for EHF hearing and age group (n=9 to 15) failed to reach statistical power for ANOVA and post-hoc comparisons, a t-test for paired data (factor hemisphere) with correction for a type−1 error (alpha shift) was applied following the Bonferroni-Holm procedure. To account for whether EHF impaired participants had smaller brain responses in either age group, descriptive information is given in the respective figures when the EHF hearing impaired group falls within or outside the 85% confidence interval (CI85%) of the EHF good hearing group (Microsoft Excel Version 2508 Build 16, Microsoft, Redmond, WA, USA). To describe the absence of an effect, 80 to 90% type−2 error values are accepted statistical guidelines for the power analysis of beta confidence level. Furthermore, to inspect for a trend across age in the different EHF hearing groups, Pearson correlation linear regression of brain responses over age was calculated. The trend was considered to be of statistical significance if the observed value of the regression coefficient **r** and group size **N** gave a probability of p<0.05. The same regression analysis was encountered for correlations between **ABR amplitudes and age or ABR amplitudes and AEFs**. ABR wave I amplitudes in categorized age groups were compared by 1-way ANOVAs for simple effects of age and EHF hearing and Holm-Sidak’s multiple comparisons test (GraphPad Prism).

In the figures and legends, statistical significance is reported for ANOVA as P-values for the respective factors and the post-hoc multiple comparisons tests and Pearson correlation momentum as asterisks, with * p<0.05, ** p<0.01, *** p<0.001. (*) indicates a trend 0.05<p<0.1. Group means falling outside the CI85% of the comparison group are indicated by ++.

## Results

### Reinforced left-hemispheric dominant AEF response to speech units, only slightly minimized by the distracting speaker, indicate higher hierarchical linguistic cortical coding

A one- or two-speaker paradigm was used to analyze frequency-tagged cortical responses from 129 MEG channels per hemisphere in response to attended and unattended, 4-word or 5-word sentences in 68 subjects of different ages as described in the methods. The topography of neural responses to the 4-word (4 Hz) or 5-word (3.75 Hz) rhythm of averaged AEF, with power spectral density in fT^2^/Hz at the sensor level, revealed strong signals in the temporal cortex in both hemispheres. Larger response signals (average power) were seen for both 4- and 5-word rhythms to one attended speaker alone (Fig. 1A, one-speaker attended) compared to the attended speaker in the presence of a second, distracting speaker (Fig. 1A, two-speaker attended, P<0.001, see also SI appendix, Table S3, 1A). For the unattended condition (Fig. 1A, two-speaker unattended), the average power declined even further (P<0.001). Power-spectral density for the control condition was close to detection level (Fig. 1A, control, P<0.001). Control conditions inspect the crosstalk between analyzed and stimulated rhythms in neuromagnetic brain responses (Fig.1 A, control, 4 Hz response to the 3.75 Hz stimulus and 3.75 Hz response to the 4 Hz stimulus). Black dots in Fig.1A indicate channels with SNR > 5.0 compared to response segments with randomly shuffled phases. Effects were statistically verified for average evoked power and topographic distribution using permutation analysis. While the topographic distributions were not different in the attended conditions (Fig. 1A, 4 Hz P=0.688, 3.75 Hz P=0.139), the topographic distribution differed between the attended and unattended conditions (Fig. 1A, 4Hz P=0.033, 3.75 Hz P=0.008). The crosstalk for the topographic distribution determined from the control condition was again significantly smaller than for the unattended condition (4 Hz P=0.003, 3.75 Hz P<0.001).

**Figure 1.**
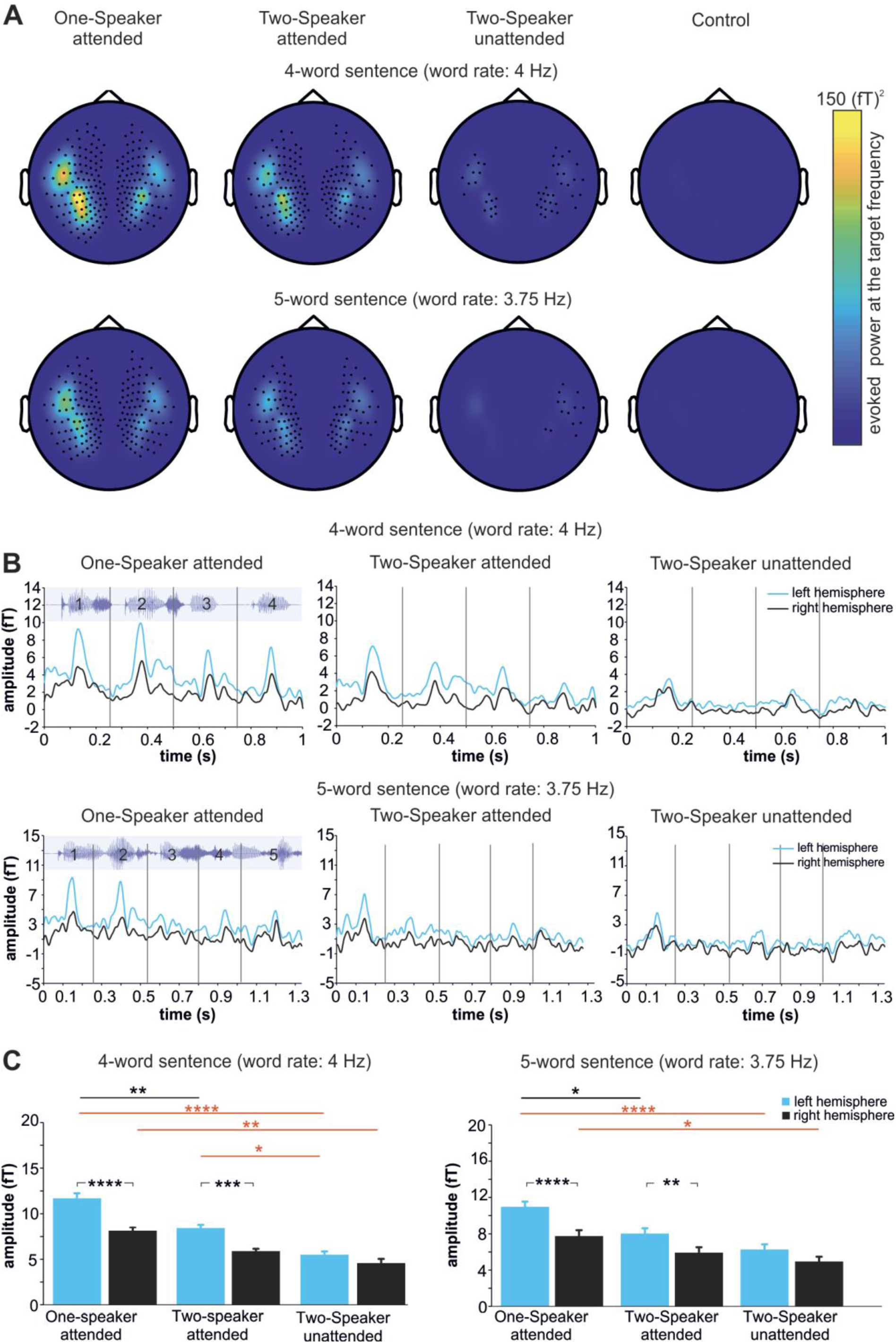
Average auditory evoked magnetic fields (AEF). (**A**) Topographic view of AEF power-spectral density in (fT)^2/Hz at the sensor level for a single attended speaker (One-speaker attended), two speakers when the target speaker is attended to (Two-speaker attended) or the unattended speaker rhythm was analyzed (Two-speaker unattended) and the crosstalk control (Control). AEF conditions were analyzed according to the 4-word and 5-word rhythm. Black dots inside the color maps indicate sensor positions at which average evoked power was at least five times larger than the expected power based on randomly shuffle phases (SNR>5.0). Attended speech led to stronger AEFs in temporal regions with left-hemisphere dominance. (**B**) Time course of the control-normalized evoked neuromagnetic response average as the absolute value of the sensor-level MEG signals in response to One-speaker attended, Two-speaker attended, and Two-speaker unattended conditions epoched for 4-word (B, upper panel) and 5-word (B, lower panel) sentences. Left hemispheric responses (blue lines) showed higher responses than right hemispheric responses (black lines), particularly under attended conditions. Numbers and waveforms in the left panels illustrate the rhythm of the voice stimuli (4-word “das Bier schmeckt gut”, 5-word “der Reis ist zu heiß”) (**C**) Quantification of the amplitude peak responses within defined time windows according to the word rhythm (numbers and vertical lines in B). A significantly higher left hemispheric response (blue bars) in attended conditions in comparison to the right hemispheric response (black bars) was observed for the attended conditions. As already shown in (A), a second speaker diminished the AEF response, with smallest amplitudes in the unattended conditions (black/red significant bars and asterisks; 3-way RM-ANOVA factor hemisphere F(1,201)=54.73 P<0.0001, factor stimulus condition F(2,201)=17.94 P<0.0001, Tukeýs multiple comparison test, * p<0.05, ** p<0.01, **** p<0.0001).

The time course of the absolute value of the neuromagnetic strength of AEF signals, normalized by subtracting the control, is shown for 4-word (Fig. 1B, upper row) and 5-word sentences (Fig.1B, lower row). Consistently higher left-hemisphere brain response amplitudes (Fig. 1B, blue lines, averaged across the 129 sensors per hemisphere) were observed compared to the right hemisphere (Fig. 1B, black lines). The left-hemisphere dominance (total-time average amplitude left-right) was, for 4-word and 5-word, 1.67 fT and 1.6 fT in the One-speaker attended condition, 1.49 fT, and 1.27 fT for the Two-speaker attended condition, and 0.68 fT and 0.85 fT for the Two-speaker unattended conditions, respectively.

As shown in Fig. 1C and the SI appendix, Table S3, 1C, when analyzing amplitude peak responses within defined time windows according to the word rhythm, the left and right hemisphere amplitude peak response was significantly different for both 4-word (Fig. 1C, left) and 5-word conditions (Fig. 1C, right) between the attended conditions (Fig. 1C, black significance bar and asterisks). This was also true between the attended conditions and the unattended condition (Fig. 1C, red significance bars and asterisks) (3-way RM-ANOVA factor hemisphere F(1,201)=54.73 P<0.0001, factor stimulus condition F(2,201)=17.94 P<0.0001). No significant effect of the word rate was found (3-way RM-ANOVA factor word rate F(1,201)=0.1467 P=0.7021). The hemisphere-specific brain activity in temporal brain regions indicates that the chosen stimulus paradigms allow for the analysis of attention-related changes in AEF responses in the cortical structures required for speech processing.

### Impaired EHF hearing lowered bilateral hemispheric cortical evoked responses during speech comprehension and led to reinforced left-hemisphere responses even in young adults

Aiming to investigate a possible role of EHF hearing on speech intelligibility during attended speech on cortical level, participants were subdivided over all ages into those with better (Fig. 2A, good EHF, blue, black) and poorer EHF hearing (Fig. 2A, impaired EHF, orange, grey), see also SI appendix, Table S1.

**Figure 2.**
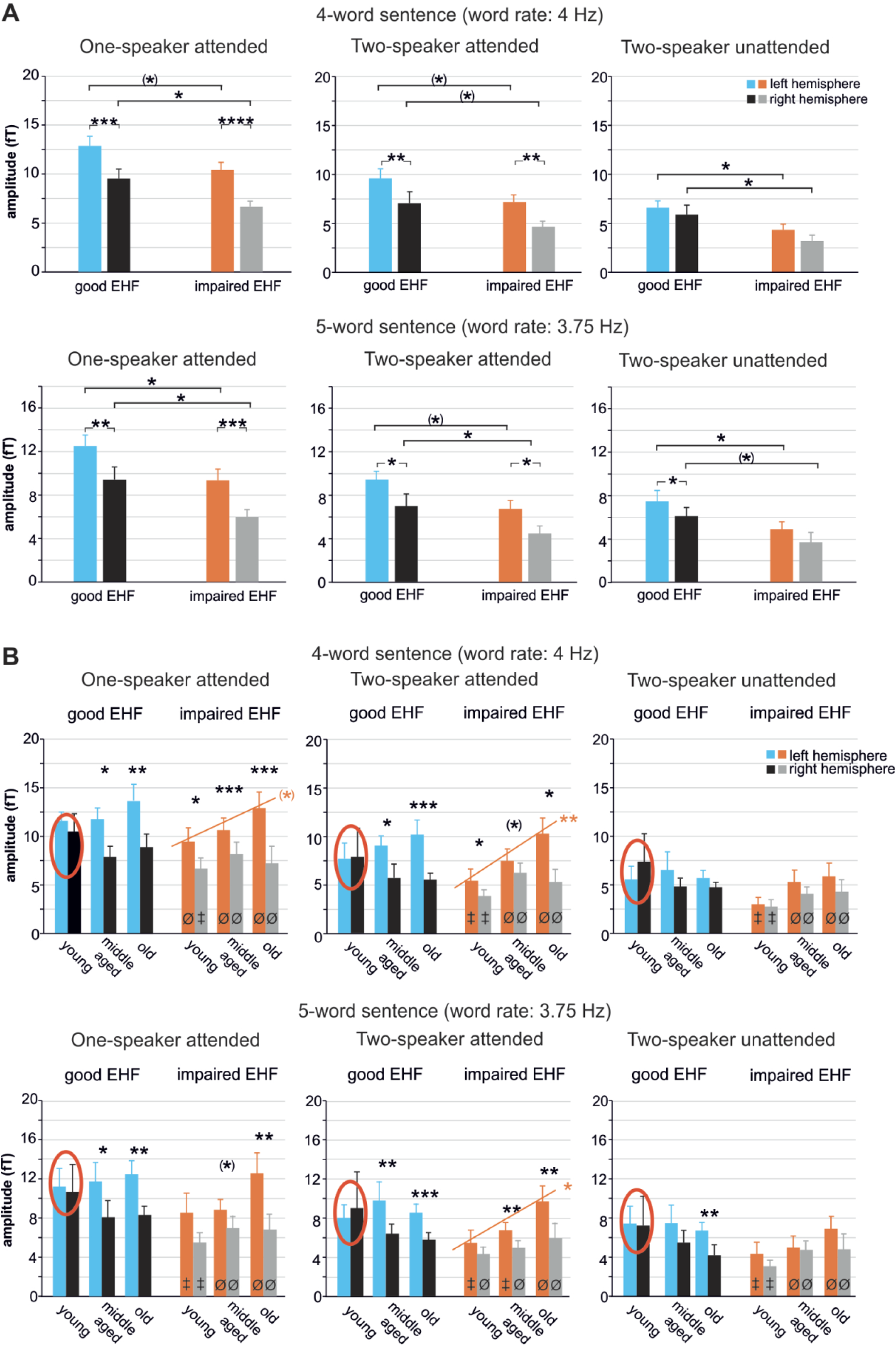
Average evoked auditory field potential (AEF) depending on EHF and age. (**A**) Overall higher left hemisphere AEF amplitudes were found for both 4-word and 5-word conditions (2-way ANOVA, factor left/right hemisphere: One-speaker attended 4-word P<0.0001, 5-word P<0.0001; Two-speaker attended 4-word P<0.0001, 5-word P=0.0006; Two-speaker unattended 4-word P=0.071, 5-word P=0.011). Impaired EHF hearing is linked with significantly reduced AEF, independently of speaker conditions and word rate. (2-way ANOVA, factor EHF good/impaired: One speaker attended 4-word P=0.0117, 5-word P=0.0039; Two-speaker attended 4-word P=0.0166, 5-word P=0.0109; Two-speaker unattended 4-word P=0.006, 5-word P=0.011). Post-hoc Sidak’s multiple comparisons test, (*) 0.05<p<0.1, * p<0.05, *** p<0.001, **** p < 0.0001. (**B**) Gradually increased left but not right hemispheric AEF amplitudes over age were found in attended conditions. A significant linear correlation over age was only observed in the attended conditions for the impaired EHF groups (orange lines and asterisks, One-speaker attended 4-word P=0.053; Two-speaker attended 4-word P=0.001, 5-word P=0.039). Paired t-test correct for multiple testing by Bonferroni-Holm, (*) 0.05<p<0.1, * p<0.05, ** p<0.01, *** p<0.001. No left hemispheric dominance (black asterisks) was observed in the young group with good EHF hearing (red circles). Symbols in the bars for the impaired EHF groups (orange, gray bars) illustrate inclusion (‡) and exclusion (Ø) in the CI85% for the good EHF groups (blue, black bars).

A reinforced left hemispheric AEF amplitude in comparison to the right hemispheric amplitude was observed for 4-word and 5-word rhythms, mainly under attended conditions (Fig. 2A, SI appendix, Table S3, 2A, hemisphere left vs right). Important for the interpretation of these findings is that a significant left hemispheric reinforcement is observed in attended speech, and is nearly independent of the presence of a distracting speaker (Fig. 2A, SI appendix, Table S3, 2A, Group comparison). AEF amplitudes were lower in all conditions in the group with impaired EHF hearing in comparison to the group with good EHF hearing (Fig. 2A, SI appendix, Table S3, 2A, Good vs impaired EHF). Group comparison using 2-way ANOVA followed by Sidak’s multiple comparisons test confirmed a statistically significant lower AEF in the group with impaired EHF, although partly only with a trend (Fig. 2A, SI appendix, Table S3, 2A, Group comparison). This indicates that impaired EHF hearing, when averaged across age, generally reduces cortical AEF amplitudes during speech processing in the right and left hemisphere.

When speech-evoked AEF responses were specifically analyzed for individual age groups (Fig. 2B, SI appendix, Table S1), a gradually increased left-hemispheric reinforcement was observed, mainly for attended conditions. Specifically for the impaired EHF group, a significant left-hemisphere reinforcement was observed to attended speech with and without distracting speaker in all age groups, but not in unattended speaker conditions (Fig. 2B black asterisks, SI appendix, Table S3, 2B, paired t-test). For the good EHF group, a similar left-hemispheric reinforcement was observed - with the exception of the young group, who had no left-hemispheric reinforcement in all conditions, strengthening the robustness of the finding (Fig. 2B red circles).

As already observed for the averaged aged groups shown in Figure 2A, the AEF amplitudes in groups with impaired EHF in comparison to good EHF were lower in each of the three different age groups. Due to the small group sizes, the statistical power was not reached to test for statistical significance. We therefore decided to test for lower AEF amplitudes by comparing the mean of the impaired EHF group with the 85% confidence interval around the good EHF group. Primarily the amplitudes of the young group with impaired EHF fell outside the confidence interval (Fig. 2B ‡).

Furthermore, to inspect for a trend over age in the different EHF hearing groups, basic linear regressions of brain-response amplitudes over age were calculated (Fig. 2B orange lines and asterisks, SI appendix, Table S3, 2B, progression over age). A significant linear correlation of left-hemisphere reinforcement and age was observed in groups with impaired EHF for almost all attended conditions, but not in unattended speaker conditions.

**Overall, the findings** so far indicate that impaired hearing in frequency regions above 8 kHz decreased cortical AEF amplitudes to attended and unattended speech in the left and right hemisphere, particularly in young adults. Bilateral lowering of AEF amplitudes to speech in participants with impaired EHF hearing occurred independently of an overall gradual increase in left-hemisphere amplitudes over age. The absence of stronger left-hemisphere AEF in attended speech in young adults with good EHF hearing suggests that intact sound energy above 8 kHz can postpone the age at which reinforcement of left hemisphere responses is required for active speech comprehension.

### Impaired EHF hearing affects reinforcement of left hemisphere AEF amplitudes through lowering stimulus onset synchrony (ABR wave I and VI)

We next reconsidered that impaired EHF hearing had previously been shown to influence synchronous auditory activity in speech-relevant frequency spectra, as shown through click-stimulus-induced ABR analysis and phoneme-induced psychometric or subcortical responses that rely on phase-locking frequency coding below 1500 Hz (14, 15). ABR wave I-VI peak amplitudes in participants with either good or impaired EHF hearing were correlated either with age (Fig. 3A), or left and right hemispheric AEF responses during attended and unattended speech (Fig. 3B, Fig. 4). A gradual decline of ABR wave I to III was observed in participants with good EHF hearing over age (Fig. 3A, blue lines, circles, blue shaded panel, SI appendix Table S3, 3A). The correlation coefficient reached significance for participants with good EHF hearing for ABR wave I, and with a statistical trend for ABR wave II and III (Fig. 3A, good EHF, ABR wave I P=0.001; ABR wave II P=0.067; ABR wave III P=0.089). The correlation coefficient of ABR wave V and VI over age did not reach statistical significance in either EHF hearing group (Fig. 3A).

**Figure 3.**
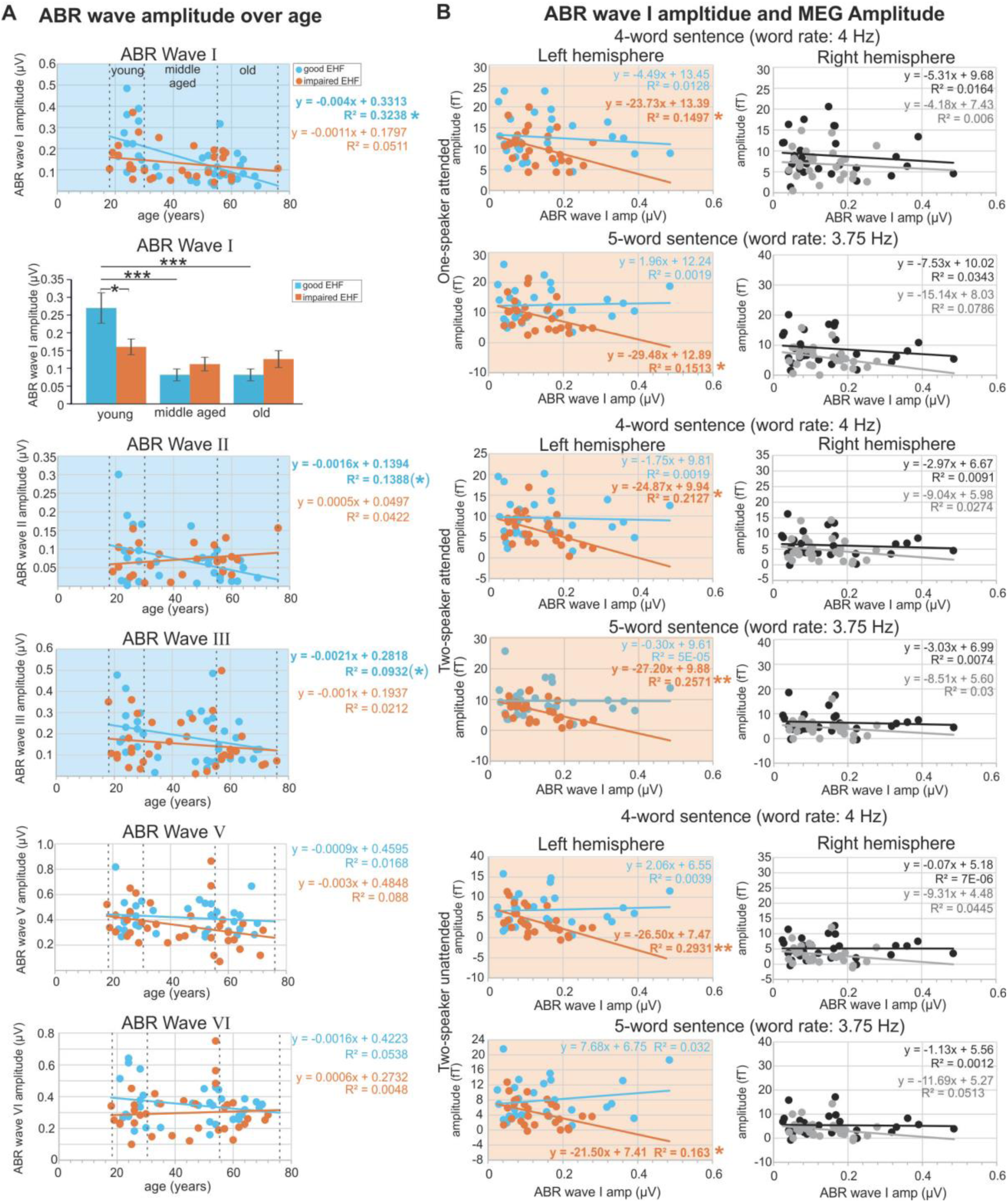
ABR wave I peak over age and in correlation to the MEG amplitude. (A) ABR wave I, II, III, V, and VI amplitudes over age. Early ABR wave amplitudes I-III decrease over age for the good EHF group (blue shaded panels, good EHF, ABR wave I P=0.001; ABR wave II P=0.067; ABR wave III P=0.089, see SI appendix Table S3 3A). Single participants - circles, linear regression with formula and correlation coefficient (*) 0.05<p<0.1, * p<0.05. The bar graph shows averaged ABR wave I amplitude data for the three age groups. A decrease over age is observed. Post-hoc Holm-Sidak’s multiple comparisons test, * p<0.05, *** p<0.001. Linear regression of AEF amplitude over ABR wave I amplitudes for the left (left column) and right hemisphere (right column). Good EHF groups in blue and black, impaired EHF group in orange and gray (left and right columns, respectively). Circles, single participants. Lines, linear regression with formula and correlation coefficient * p<0.05, ** p< 0.01. Panels in color indicate significant correlations, or a statistical trend of the respective group between ABR wave I amplitude and AEF amplitude.

**Figure 4.**
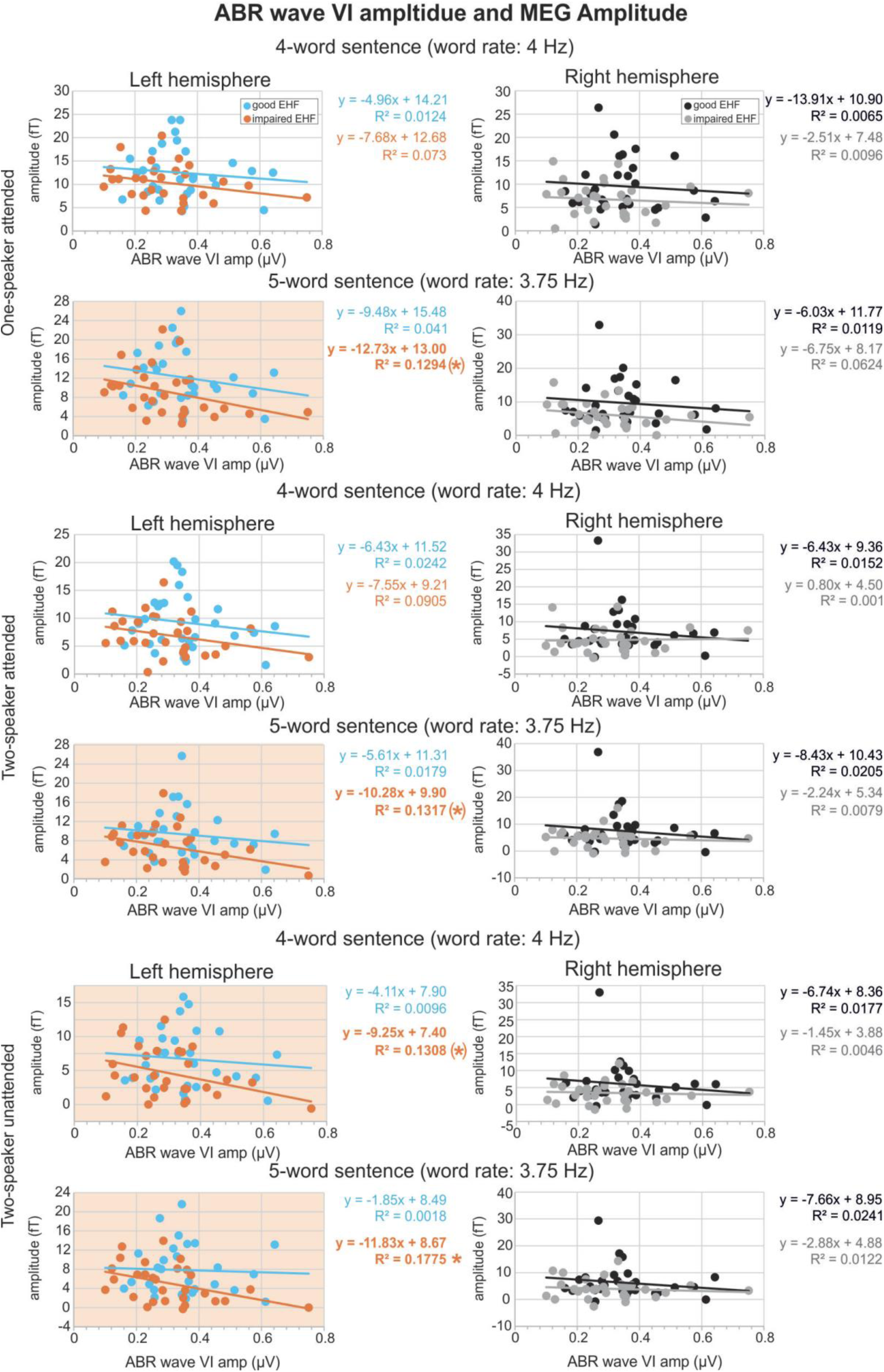
ABR wave VI peak in correlation to the MEG amplitude. Linear regression of AEF amplitude over ABR wave VI amplitudes for left hemisphere (left column) and right hemisphere (right column). Good EHF groups in blue and black, impaired EHF group in orange and gray (left and right columns respectively). Circles, single participants. Lines, linear regression with formula and correlation coefficient (*) 0.05<p<0.1, * p<0.05. The lower values and regression line for the EHF impaired group illustrate smaller neuromagnetic AEF amplitudes. Panels in color indicate significant correlations, or a statistical trend of the respective group between ABR wave VI amplitude and AEF amplitude.

The significant decrease of ABR wave I amplitude over age for the group with good EHF hearing was confirmed by averaging the data for the age groups defined in SI appendix Table S2 (Fig.3A, bar graph, blue bars, black asterisks, 1-way ANOVA P<0.0001, F(5, 48) = 794, Holms Sidak’s multiple comparisons test young vs. middle-aged P<0.0001, young vs. old P<0.0001). No age progression was found for the impaired EHF group (Fig.3A, orange bars), and a significant difference between the good and the impaired hearing groups was only found for young adults (Fig. 3A, bar graph, young, Holms Sidak’s multiple comparisons test P=0.014). The significant difference in the regression slope of ABR wave I between good and impaired EHF hearing (Fig. 3A) indicates that good EHF hearing, in which reinforced left-hemisphere dominance of AEF has not yet occurred (Fig. 2B, blue bar), exhibits higher ABR wave I peak amplitudes, while the group of young adults with impaired EHF hearing in which reinforced left-hemisphere dominance of AEF had already occurred (Fig. 2B, orange bar), exhibited lower ABR wave I amplitudes (Fig. 3A). ABR wave I amplitudes were not correlated with AEF amplitudes of the **right** hemispheres (Fig. 3B, right column, SI appendix Table S3, 3B). For the left hemisphere AEF, amplitudes correlated with ABR wave I amplitudes for impaired EHF groups for all conditions (Fig. 3B, left column, orange regression lines and circles, orange shaded panels, impaired EHF; One speaker attended 4-word P=0.046, 5-word P=0.045; Two speaker attended 4-word P=0.016, 5-word P=0.007; Two speaker unattended 4-word P=0.004, 5-word P=0.037). This shows that participants with impaired EHF hearing exhibited smaller left-hemispheric amplitudes, even when ABR wave I amplitudes were high. This also indicates that a balanced cortical neuromagnetic AEF response requires good EHF hearing in functional high frequency auditory nerve responses.

More or less the same is found when AEF amplitudes in subjects with good or impaired EHF conditions were correlated to ABR wave VI peak amplitudes (Fig. 4, SI appendix Table S4, 4). Here again, higher amplitudes were measured in participants with good EHF hearing (Fig. 4A, blue and black symbols and lines) as compared to impaired EHF hearing (Fig. 4, orange and gray symbols and lines). AEF amplitudes in the left hemisphere correlated with ABR wave VI amplitudes for One speaker attended 5-word (Fig. 4 left column, orange shaded panels, P=0.055), Two speaker attended 5-word (Fig. 4 left column, P=0.053) and Two speaker unattended 4-word (Fig. 4 left column P=0.054) and 5-word (Fig. 4 left column P=0.023).

As also shown for cortical response relations to ABR wave I, the difference between good and impaired EHF hearing remained relatively small when AEF amplitudes were higher (Fig. 2B) and ABR wave VI was small in size (Fig. 4), as suggested for older subjects (14). On the other hand, when ABR wave VI amplitudes were larger (likely younger or middle aged subjects, (14)), subjects with impaired EHF can also only increase AEF amplitudes to a limited extent (Fig. 4).

## Discussion

While the influence of extended high-frequency (EHF) deficits on speech comprehension in noise has been discussed in previous studies, the present study provides three main new contributions: **First**, we showed evidence that sound energy above 8 kHz (EHF hearing) contributes to cortical activity during the processing of higher hierarchical linguistic units. **Second**, we found that impaired EHF hearing accelerates the onset of the reinforcement of left-hemisphere cortical activity during speech intelligibility across age. **Third,** we demonstrate that EHF hearing influences neural cortical activity during reinforced left-hemisphere activity via its impact on synchronized auditory responses at stimulus onset. These aspects are discussed in the following.

### EHF hearing influences evoked cortical activity during higher hierarchical speech processing

Various studies have suggested a role of EHF hearing > 8kHz for speech recognition and speech perception in noise (See for a review (11, 12, 20, 21)). By subdividing subjects of different ages into those with good or impaired EHF hearing, we here demonstrate for the first time a profound decline of the overall auditory evoked cortical response during speech comprehension and intelligibility in subjects with impaired versus preserved EHF hearing (Fig. 2A, SI appendix, S2). We assume that this indicates that sound energy at frequencies above 8 kHz critically influences the processing of higher order linguistic structures for the following reasons: **(i)** The altered evoked auditory field potential observed in response to attended or unattended 4- or 5-word rate speech was tagged within slow oscillation frequencies of 4 Hz or 3.75 Hz. This frequency range falls within the theta band (3-7 Hz) in which cerebro-acoustic phase locking particular during speech intelligibility, under attentional modulation, exhibits its highest power (6, 7). **(ii)** The evoked AEF changes in response to speech units were localized bilaterally to the temporal lobe (Fig. 1A), as expected from auditory evoked cortical field responses (16, 22). **(iii)** Speech processing in the human brain engages both cerebral hemispheres that contribute to processing different aspects of speech (23, 24). The right hemisphere plays a dominant role in the processing of suprasegmental features of speech, including e.g., pitch or prosody segments (24–27). On the other hand, speech processing at highest hierarchical levels requires fine-grained temporal resolution and, when under the influence of attention, is modulated by activity from the medial temporal gyrus and the hippocampus. This process is assumed to reinforce a left-hemispheric dominance (5, 7, 24, 28–35).

The process underlying this left-lateralization shift is suggested to occur when bilateral or right-hemispheric auditory cortical activity is modulated by linguistic information from non-primary sensory regions, specifically within the temporal lobe that is involved in attention, leading to increased synchronization of the envelope of speech (17, 18, 36–38). During this process phase locking of ongoing neural oscillation is increased, particularly the slow theta oscillation of 4-8 Hz rate, (6, 39). Impaired EHF hearing was associated with reduced AEF amplitudes in response to attended or unattended 4- and 5-word rate tagged with 4 Hz and 3.75 Hz, respectively, and was so similarly in the right and left hemispheres (Fig. 2A, SI appendix, Tables S2). This suggests that sound energy above 8kHz improves both the integration of auditory and non-auditory cues towards left-hemispheric reinforcement and the necessary pre-processing cortical steps for this. **(iv)** Moreover, the influence of EHF hearing on higher hierarchical cortical processing was confirmed by the observation that impaired EHF hearing reduced evoked AEF amplitudes irrespective of a second distracting speaker (Fig. 2A,B, SI appendix, Tables S2). As such, it is a characteristic of left-hemispheric response increase to speech signals of higher hierarchical linguistic structures that the cortical activity becomes less reduced by distracting speakers under attended conditions in comparison to, e.g., speech signals of lower hierarchical linguistic structure (6, 18, 32, 33, 39–46). This likely reflects the contribution of memory-related processes, required for hierarchical structure-building operations during language comprehension, involve an automatic re-activation or ‘retrieval’ of memory traces. These processes place reduced demands on subcortical auditory resources and therefore less affected by distracting auditory input (33, 39, 47, 48).

### EHF hearing, when impaired, accelerates the onset of left-hemisphere reinforcement during speech intelligibility across age

We here observed a clear age-dependent increase in left-hemispheric amplitudes independent of preserved or impaired EHF hearing, mainly in attended speech (Fig. 2B). A striking absence of this reinforced left-hemispheric AEF amplitude was seen in younger participants (19-28 years) with preserved EHF hearing (Fig. 2A,B, SI appendix, Tables S2). This finding emphasizes that the left-hemisphere reinforcement of cortical amplitudes across age is possibly accelerated to an earlier age in the case that EHF hearing is impaired. Several previous studies have observed reinforced cortical responses over aging, not only for left-, but also for right-hemispheric associated auditory processing (26, 49–55).

The functional significance of enhanced cortical responses in older age is not well understood. Although the higher cortical amplitude over age may apparently be of advantage, as it may ensure sustained attention in an impoverished acoustic environment, as occurs over age, it does so at the cost of limited cognitive resources (26). Thus, increasing evidence that enhanced cortical activity across age, independent of affecting left or right hemispheres, is associated with poorer performance in a test of cognition, requiring more effort in listening (26). Consistent with this, an increased laterality index in the elderly has been proposed to reflect greater by difficulty in maintaining attentional focus (56). These are accompanied by deficits in temporal processing (57), or by deficits in pitch perception based on periodicity (58), with diminished neural synchronization to speech related acoustic modulations (50) or smaller FFRs to steady vowel sounds being reported (59). Mechanistically, age-dependent changes in lateralization have been suggested to reflect a decline in interhemispheric inhibitory control (e.g. (49, 51)).

It can therefore be concluded that avoiding a compensatory increase of cortical amplitudes over age – as shown here for left-hemispheric cortical AEF responses in subjects with good EHF hearing, is beneficial. The additive use of cortical cognitive resources could be avoided, and optimized speech comprehension functions could be preserved.

### EHF hearing influences neural cortical activity through its impact on synchronized auditory responses at the stimulus onset

We demonstrate here that impaired EHF hearing can be associated with diminished ABR wave I amplitudes (Fig. 3A), even in younger individuals (Fig. 3A). Notably, even at the youngest age (here 19-29 years), this was accompanied by an increase of the left-hemispheric AEF amplitude (Fig. 2B), indicating a direct link between ABR wave I size and left-hemispheric reinforcement. When the deterioration of higher-frequency hearing occurs later in life, the cortical AEF amplitude remains reduced, despite higher ABR wave I and VI amplitudes (Fig. 3, 4; SI appendix, Tables S3, 3,4), highlighting the influence of sound energy above 8 kHz for maintaining appropriate cortical response amplitudes. Extended high-frequencies were previously shown to impact synchronous phase-locked activity at lower stimulation frequencies (60), as well as ABR wave I peak amplitudes through their impact on high-SR auditory nerve fibers **(ANF**) (14, 15).

Through their influence on synchronized responses high spontaneous-rate (SR) ANFs, which have low activation thresholds, primarily determine ABR wave peak amplitudes at stimulus onset (61). In contrast, low spontaneous-rate ANFs, characterized by higher activation thresholds, do not contribute to the synchronicity of ANF (61). Also, outer hair-cell function could be excluded as the factor that, instead of EHF frequencies, influenced temporal coding and phase-locking of ANF in response to frequency-following responses (60). In support of EHF hearing exhibiting an impact on speech perception on synchronization, is the observation that EHF hearing was linked to a self-reported difficulty of hearing in a noisy environment (11). The self-reported evaluation of speech-comprehension ability was previously shown to be better explained by changes in synchronized responses at the stimulus onset, rather than by hearing loss across age (14). Numerous excellent reviews collecting factors contributing to age-related hearing loss in brain and cognitive function (9, 10, 62) might – based on the present study-add EHF hearing as a critical contributor to higher hierarchical speech processing over age.

### Conclusion

We suggest that sound energy above 8kHz promotes well-synchronized auditory responses at the stimulus onset, and thereby improves the process of proper phase locking of auditory and non-auditory cues during speech processing. This implies that the successful maintenance of speech intelligibility over age, which has been proposed to depend on limited cognitive resources, as well as on working memory capacity or microvascular resources (5, 9, 10, 63, 64), may benefit from preserved EHF hearing. The present findings therefore bear directly on the question of whether early EHF hearing loss may act as a precursor of more severe hearing decline in later life (11). Regarding the limitation of cognitive resources as a risk factor in dementia (64), this leads to the most intriguing question as to whether earlier prevention of hearing loss in EHF ranges by means of e.g. earlier restoration of EHF hearing using hearing aids (if technically feasible) would lower the risk of losing cognitive performance with age? This perspective may motivate the development of technologies that address current limitations in clinical audiology and hearing-aid fitting, particularly by extending coverage to historically neglected frequency range above 8 kHz in hearing.

## Abbreviations

**ABR:** auditory brainstem responses, **AEF:** auditory evoked cortical fields, **ANF**: auditory nerve fiber, **ANOVA**: one-way analysis of variance, **EEG:** electroencephalography, **EHF:** extended high frequencies, **FFR:** frequency-following responses, **ICA:** independent component analysis, **MEG:** magnetoencephalography, **PTA-EHF:** pure-tone average for extended high frequencies, **SNR:** signal-to-noise ratio, **SR:** spontaneous rate.

## Data and code availability statements

The raw data supporting the conclusions of this article will be made available by the authors on request.

## Acknowledgments

This work was supported by the Deutsche Forschungsgemeinschaft DFG KN 316/13-1, DFG RU 713/6-1; ERA-NET NEURON JTC 2020: BMBF 01EW2102 CoSySpeech and FWO G0H6420N; the IZKF Promotionskolleg of the Faculty of Medicine, University Hospital of Tübingen. VICI Grant (Grant No. 918-17-603); the Netherlands Organization for Scientific Research (NWO), and the Netherlands Organization for Health Research and Development (ZonMw); CherISH, European Union’s Horizon research and innovation programme under Marie Sklodowska-Curie Grant Agreement No: 101120054. The Spanish PID2023-148541OB-I00, funded by MICIU/AEI https://doi.org/10.13039/501100011033 and FEDER EU; the Consejería de Educación, Junta de Castilla y León (SA218P23), and the strategic research programs of excellence from the Regional Government of Castile and León, co-funded by the ERDF Operational Programme (ref. CLU-2023-1-01). Programa de Doctorado en Neurociencias, Centro Interdisciplinario de Neurociencia UC, Pontificia Universidad Católica de Chile, Santiago, Chile. The funding source(s) were not involved in study design, data collection, analysis or interpretation, or in writing and submitting the manuscript.

We thank the audiometrists of the HNO-Klinik Tübingen for their expertise in recording audiograms, and stels-ol for English language services

## Author contributions

Rodrigo Dononso-San Martin: Data curation; Formal analysis; writing

Konrad Dapper: Data collection, Data curation, Formal analysis, Investigation Software,

Stefan Fink: Data curation; Formal analysis;

Etienne Gaudrain: Methodology, Software;

Deniz Başkent: Methodology, Software;

Sarah Verhulst: Methodology, Software; Editing;

Csaba Harasztosi: Formal analysis

Wibke Singer: Writing, Editing

Manuel Malmierca: Edting of the MS Markus Siegel: Methodology, Editing;

Ernst Dalhoff: Methodology;

<u>Stephan M. Wolpert:</u> Methodology, Supervision;

Christoph Braun: Project administration Methodology Supervision, Writing - original draft Writing - review & editing

Marlies Knipper & Lukas Rüttiger: Funding acquisition Methodology, Statistics, Project administration, Supervision, Writing - original draft/ Writing - review & editing;

