## Supplement Material for "Extended high frequency hearing influences cortical response amplitudes to speech"

**Donoso San-Martin et al**

Supplementary Information

### Figures

### Supplementary Figure 1

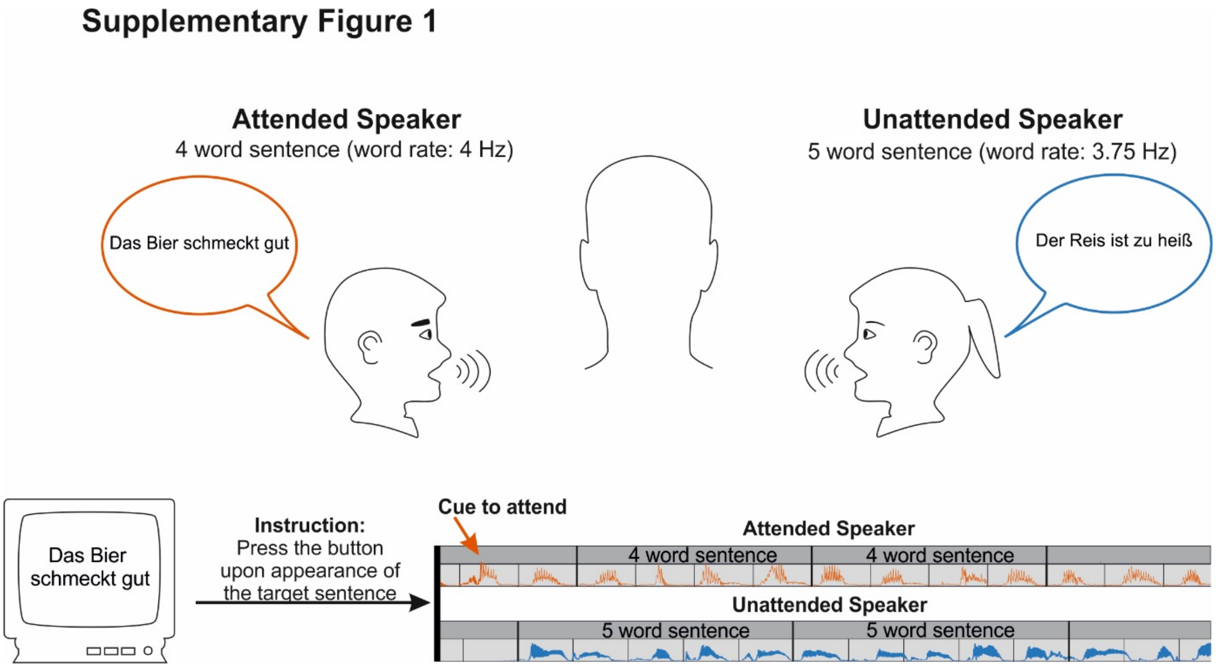


**Supplementary Figure 1***: Design of the speech-comprehension task during the MEG recordings. One- or two-speakers with different voices, speaking rate (word rate and sentence rate) present meaningful, easy sentences of 4- or 5-words. 4-word sentences had a word rate of 4 Hz. 5-word sentences had a word rate of 3.75 Hz. The task was to detect the target sentence (visually cued on the screen) spoken via the "Attended Speaker" with or without a competing second speaker ("Unattended Speaker"). Subjects were instructed to pay attention to the Attended Speaker, the one that speaks first and randomly says the target sentence.*

**Tables**

Table S1: Participant groups and numbers

| *Participants included* | N = 68 | | |
|---|---|---|---|
| *Audiometry (PTA-EHF)* | | | |
| | participants tested | EHF good | EHF impaired |
| All | 68 | 35 | 33 |
| Young (18-29 years) | | 10 | 9 |
| Middle aged (30-55 years) | | 10 | 14 |
| Old (56-76 years) | | 15 | 10 |

***Table S1:*** *Number of participants included in the study. Of 81 people measured in the MEG, 3 were ruled out following the measurement because of metal implants (e.g., dental braces), 3 after technical problems arose, 4 due to non-compliant behavior during the examination (misunderstood the task or fell asleep), and 3 as high-frequency audiometry data were not available. From the remaining 68 participants, all data were included.*

Table S2: Stimulus conditions

| Conditions 3 x 40 block, 29 x 4 word, 23 x 5 word | |
|---|---|
| Block Ia | |
| | •Target Speaker 1 left (4 Hz) |
| | •Target Speaker 1 left (3.75 Hz) |
| | •Target Speaker 2 right (4 Hz) |
| | •Target Speaker 2 right (3.75 Hz) |
| Block Ib | |
| | •Target Speaker 1 left (4 Hz) and Distractor Speaker 2 right (3.75 Hz) |
| | •Target Speaker 1 left (3.75 Hz) and Distractor Speaker 2 right (4 Hz) |
| | •Target Speaker 2 right (4 Hz) and Distractor Speaker 1 left (3.75 Hz) |
| | •Target Speaker 2 right (3.75 Hz) and Distractor Speaker 1 left (4 Hz) |
| Block IIa | |
| | •Target Speaker 1 left and right (4 Hz) |
| | •Target Speaker 1 left and right (3.75 Hz) |
| | •Target Speaker 2 right (4 Hz) |
| | •Target Speaker 2 right (3.75 Hz) |
| Block IIb | |
| | •Target Speaker 1 left and right (4 Hz) and Distractor Speaker 2 right (3.75 Hz) |
| | •Target Speaker 1 left and right (3.75 Hz) and Distractor Speaker 2 right (4 Hz) |
| | •Target Speaker 2 right (4 Hz) and Distractor Speaker 1 left and right (3.75 Hz) |
| | •Target Speaker 2 right (3.75 Hz) and Distractor Speaker 1 left and right (4 Hz) |
| Block IIIa | |
| | •Target Speaker 1 left and right (4 Hz) |
| | •Target Speaker 1 left and right (3.75 Hz) |
| | •Target Speaker 2 left and right (4 Hz) |
| | •Target Speaker 2 left and right (3.75 Hz) |
| Block IIIb | |
| | •Target Speaker 1 left and right (4 Hz) and Distractor Speaker 2 left and right (3.75 Hz) |
| | •Target Speaker 1 left and right (3.75 Hz) and Distractor Speaker 2 left and right (4 Hz) |
| | •Target Speaker 2 left and right (4 Hz) and Distractor Speaker 2 left and right (3.75 Hz) |
| | •Target Speaker 2 left and right (3.75 Hz) and Distractor Speaker 2 left and right (4 Hz) |

***Table S2:*** *Outline of the recording session in 3 blocks (I to III), each containing 40 segments of 30 seconds. Within each 30s segment, 29 4-word sentences, or 23 5-word sentences, were presented, either via one speaker (only Target Speaker) or both speakers together (the target speaker to be attended to, and the distractor speaker to be disregarded). In Blocks Ia to IIIa only the target speaker was presented, either monaural to the left or right ear or to both ears. In Blocks Ib to IIIb two speakers were active, with variation* *of speaker voices, stimulated ears, and word rates, to keep the participants attentive to the task. In each block, 4-word and 5-word sentences were included***.**

Table S3: P-values for statistical group comparisons

| Figure | Stimulus condition comparison | Analyzed rhythm | P-value<br>Avg evoked power | Topographic distribution |
|---|---|---|---|---|
| **1A** | One-speaker attended vs. Two-speaker attended | 4 Hz word rate<br>3.75 Hz word rate | < 0.001<br>< 0.001 | 0.688<br>0.139 |
| | Two-speaker attended vs. Two-speaker unattended | 4 Hz word rate<br>3.75 Hz word rate | < 0.001<br>< 0.001 | 0.033<br>0.008 |
| | Two-speaker unattended vs. Control | 4 Hz word rate<br>3.75 Hz word rate | < 0.001<br>< 0.001 | 0.003<br>< 0.001 |
| **1C** | **Condition analyzed** | | **P-value**<br>**3-way ANOVA** | **F (DFn, DFd)** |
| | Stimulus condition | | < 0.0001 | F (2, 201) = 17.94 |
| | Hemisphere | | < 0.0001 | F (1, 201) = 54.73 |
| | Word rate | | 0.7021 | F (1, 201) = 0.1467 |
| | **Stimulus conditions compared** | **Analyzed rhythm** | **Tukey's multiple comparisons test P-value**<br>**Left** | **Right** |
| | One-speaker attended vs. Two-speaker attended | 4 Hz word rate<br>3.75 Hz word rate | 0.006<br>0.026 | 0.230<br>0.588 |
| | Two-speaker attended vs. Two-speaker unattended | 4 Hz word rate<br>3.75 Hz word rate | 0.024<br>0.578 | 0.922<br>0.987 |
| | One-speaker attended vs. Two-speaker unattended | 4 Hz word rate<br>3.75 Hz word rate | < 0.001<br>< 0.001 | 0.001<br>0.038 |
| | **Stimulus condition comparison** | **Analyzed rhythm** | **P-value**<br>**Left-right** | |
| | One-speaker attended | 4 Hz word rate<br>3.75 Hz word rate | < 0.0001<br>< 0.0001 | |
| | Two-speaker attended | 4 Hz word rate<br>3.75 Hz word rate | 0.0004<br>0.009 | |
| | Two-speaker unattended | 4 Hz word rate<br>3.75 Hz word rate | 0.889<br>0.416 | |
| **2A** | **Condition analyzed** | | **P-value**<br>**2-way ANOVA** | **F (DFn, DFd)** |
| | One-speaker attended<br>Left/right hemisphere | 4 Hz word rate<br>3.75 Hz word rate | < 0.0001<br>< 0.0001 | F (1, 66) = 40.85<br>F (1, 66) = 26.80 |
| | One-speaker attended | 4 Hz word rate | 0.0117 | F (1, 66) = 6.729 |

| | | | | |
|---|---|---|---|---|
| | Good/impaired EHF | 3.75 Hz word rate | 0.0039 | F (1, 66) = 8.972 |
| | Two-speaker attended<br>Left/right hemisphere | 4 Hz word rate<br>3.75 Hz word rate | < 0.0001<br>0.0006 | F (1, 66) = 20.15<br>F (1, 66) = 12.99 |
| | Two-speaker attended<br>Good/impaired EHF | 4 Hz word rate<br>3.75 Hz word rate | 0.0166<br>0.0109 | F (1, 66) = 6.040<br>F (1, 66) = 6.861 |
| | Two-speaker unattended<br>Left/right hemisphere | 4 Hz word rate<br>3.75 Hz word rate | 0.071 (n.s.)<br>0.011 | F (1, 66) = 3.390<br>F (1, 66) = 6.886 |
| | Two-speaker unattended<br>Good/impaired EHF | 4 Hz word rate<br>3.75 Hz word rate | 0.006<br>0.011 | F (1, 66) = 8.145<br>F (1, 66) = 6.836 |
| | | | **Sidak's multiple comparisons test** | |
| | **Groups compared** | **Analyzed rhythm** | **P-value**<br>Left | **P-value**<br>Right |
| | One-speaker attended<br>Good vs. impaired | 4 Hz word rate<br>3.75 Hz word rate | 0.073 (n.s.)<br>0.026 | 0.03<br>0.016 |
| | Two-speaker attended<br>Good vs. impaired | 4 Hz word rate<br>3.75 Hz word rate | 0.063 (n.s.)<br>0.065 (n.s.) | 0.07 (n.s.)<br>0.037 |
| | Two-speaker unattended<br>Good vs. impaired | 4 Hz word rate<br>3.75 Hz word rate | 0.049<br>0.021 | 0.016<br>0.08 (n.s.) |
| | | | **Sidak's multiple comparisons test** | |
| | **Condition analyzed** | **Analyzed rhythm** | **P-value**<br>Left-right<br>Good EHF | **P-value**<br>Left-right<br>Impaired EHF |
| | One-speaker attended | 4 Hz word rate<br>3.75 Hz word rate | 0.0001<br>0.001 | <0.001<br>0.001 |
| | Two-speaker attended | 4 Hz word rate<br>3.75 Hz word rate | 0.004<br>0.036 | 0.005<br>0.019 |
| | Two-speaker unattended | 4 Hz word rate<br>3.75 Hz word rate | 0.526<br>0.049 | 0.225<br>0.293 |
| **2B** | | **Paired t-tests, P value adjusted by Bonferoni-Holms** | | |
| | **Condition analyzed** | **Analyzed rhythm** | **P-value**<br>Left-right<br>Good EHF | **P-value**<br>Left-right<br>Impaired EHF |
| | One-speaker attended | 4 Hz word rate<br>3.75 Hz word rate | 0.600<br>0.843 | 0.029<br>0.14 |
| Young | Two-speaker attended | 4 Hz word rate<br>3.75 Hz word rate | 0.848<br>0.778 | 0.066<br>0.178 |
| | Two-speaker unattended | 4 Hz word rate | 0.614 | 0.849 |

| | | | | |
|---|---|---|---|---|
| | | 3.75 Hz word rate | 0.973 | 0.49 |
| Middle-aged | One-speaker attended | 4 Hz word rate | 0.03 | 0.003 |
| | | 3.75 Hz word rate | 0.022 | 0.072 (n.s.) |
| | Two-speaker attended | 4 Hz word rate | 0.06 | 0.099 (n.s.) |
| | | 3.75 Hz word rate | 0.012 | 0.024 |
| | Two-speaker unattended | 4 Hz word rate | 0.164 | 0.102 |
| | | 3.75 Hz word rate | 0.104 | 0.704 |
| Old | One-speaker attended | 4 Hz word rate | 0.018 | 0.004 |
| | | 3.75 Hz word rate | 0.003 | 0.021 |
| | Two-speaker attended | 4 Hz word rate | 0.003 | 0.054 |
| | | 3.75 Hz word rate | 0.003 | 0.038 |
| | Two-speaker unattended | 4 Hz word rate | 0.147 | 0.148 |
| | | 3.75 Hz word rate | 0.003 | 0.288 |
| | **Progression slope over age** | | **P-value** Left | **P-value** Right |
| | One-speaker attended good EHF | 4 Hz word rate | 0.270 | 0.728 |
| | | 3.75 Hz word rate | 0.554 | 0.396 |
| | Impaired EHF | 4 Hz word rate | 0.053 (n.s.) | 0.6 |
| | | 3.75 Hz word rate | 0.132 | 0.541 |
| | Two-speaker attended good EHF | 4 Hz word rate | 0.153 | 0.430 |
| | | 3.75 Hz word rate | 0505 | 0.287 |
| | Impaired EHF | 4 Hz word rate | 0.001 | 0.304 |
| | | 3.75 Hz word rate | 0.039 | 0.398 |
| | Two-speaker unattended good EHF | 4 Hz word rate | 0.799 | 0.345 |
| | | 3.75 Hz word rate | 0.884 | 0.262 |
| | Impaired EHF | 4 Hz word rate | 0.206 | 0.508 |
| | | 3.75 Hz word rate | 0.413 | 0.506 |

| **3A** | **Progression slope over age** | **P-value** Good EHF | **P-value** Impaired EHF |
|---|---|---|---|
| | ABR wave I | 0.001 | 0.2568 |
| | ABR wave II | 0.067 (n.s.) | 0.377 |
| | ABR wave III | 0.089 (n.s.) | 0.435 |
| | ABR wave V | 0.495 | 0.112 |
| | ABR wave VI | 0.226 | 0.722 |

| | | | | |
|---|---|---|---|---|
| | **Mean ABR wave Amplitude** | | **P-value** 1-way ANOVA | **F (DFn, DFd)** |
| | ABR wave I | | < 0.0001 | F (5, 48) = 794 |
| | | | Holm-Sidak's multiple comparisons test | |
| | ABR wave I | | **P-value** Good EHF | **P-value** Impaired EHF |
| | Young vs. middle-aged | | < 0.0001 | 0.408 |
| | Young vs. old | | < 0.0001 | 0.643 |
| | Middle-aged vs. old | | >0.9999 | 0.709 |
| | | | **P-value** Good vs. impaired EHF | |
| | Young | | 0.014 | |
| | Middle-aged | | 0.470 | |
| | Old | | 0.470 | |
| **3B** | **Regression slope over ABR wave I amplitude** | **Analyzed rhythm** | **P-value** Left | **P-value** Right |
| | One-speaker attended good EHF | 4 Hz word rate | 0.560 | 0.508 |
| | | 3.75 Hz word rate | 0.821 | 0.336 |
| | Impaired EHF | 4 Hz word rate | 0.046 | 0.701 |
| | | 3.75 Hz word rate | 0.045 | 0.157 |
| | Two-speaker attended good EHF | 4 Hz word rate | 0.823 | 0.623 |
| | | 3.75 Hz word rate | 0.970 | 0.658 |
| | Impaired EHF | 4 Hz word rate | 0.016 | 0.401 |
| | | 3.75 Hz word rate | 0.007 | 0.3873 |
| | Two-speaker unattended good EHF | 4 Hz word rate | 0.749 | 0.989 |
| | | 3.75 Hz word rate | 0.353 | 0.861 |
| | Impaired EHF | 4 Hz word rate | 0.004 | 0.291 |
| | | 3.75 Hz word rate | 0.037 | 0.256 |
| **4** | **Regression slope over ABR wave VI amplitude** | **Analyzed rhythm** | **P-value** Left | **P-value** Right |
| | One-speaker attended Good EHF | 4 Hz word rate | 0.566 | 0.677 |
| | | 3.75 Hz word rate | 0.292 | 0.573 |
| | Impaired EHF | 4 Hz word rate | 0.156 | 0.613 |
| | | 3.75 Hz word rate | 0.055 (n.s.) | 0.191 |
| | Two-speaker attended good EHF | 4 Hz word rate | 0.421 | 0.524 |
| | | 3.75 Hz word rate | 0.489 | 0.459 |

| | | | |
|---|---|---|---|
| Impaired EHF | 4 Hz word rate | 0.113 | 0.871 |
| | 3.75 Hz word rate | 0.053 (n.s.) | 0.646 |
| Two-speaker unattended good EHF | 4 Hz word rate | 0.614 | 0.492 |
| | 3.75 Hz word rate | 0.826 | 0.421 |
| Impaired EHF | 4 Hz word rate | 0.054 (n.s.) | 0.727 |
| | 3.75 Hz word rate | 0.023 | 0.569 |
